# Ketamine increases activity of a fronto-striatal projection that regulates compulsive behavior

**DOI:** 10.1101/2020.07.06.190553

**Authors:** Gwynne L. Davis, Adelaide R. Minerva, Argentina Lario, Carolyn I. Rodriguez, Lisa A. Gunaydin

## Abstract

Obsessive-Compulsive Disorder (OCD), characterized by intrusive thoughts (obsessions) and repetitive behaviors (compulsions), is associated with dysfunction in fronto-striatal circuits. There are currently no fast-acting pharmacological treatments for OCD. However, recent clinical studies demonstrated that an intravenous infusion of ketamine rapidly reduces OCD symptoms. To probe mechanisms underlying ketamine’s therapeutic effect on OCD-like behaviors, we used the SAPAP3 knockout (KO) mouse model of compulsive grooming. Here we recapitulate the fast-acting therapeutic effect of ketamine on compulsive behavior, and show that ketamine increases activity of dorsomedial prefrontal neurons projecting to the dorsomedial striatum in KO mice. Optogenetically mimicking this increase in fronto-striatal activity rescued compulsive grooming behavior in KO mice. Conversely, inhibiting this circuit in wild-type mice increased grooming. These studies demonstrate that ketamine increases activity in a fronto-striatal circuit that causally controls compulsive grooming behavior, suggesting this circuit may be important for ketamine’s therapeutic effects in OCD.

## Introduction

Obsessive-compulsive disorder (OCD) is characterized by recurring, intrusive thoughts (obsessions) and repetitive behaviors (compulsions)^1, 2^. Available pharmacological treatments (serotonin reuptake inhibitors or SRIs) rarely produce complete symptom remission and take 2-3 months to produce meaningful symptom relief^3^. Often high SRI doses are required with increased side effect burden, and ~30-40% of OCD patients remain refractory to these treatments^4, 5^. Identifying effective, fast-acting treatments will help reduce OCD morbidity.

Increasing evidence indicates that disruptions in glutamate signaling may play a role in OCD symptoms^6, 7^. Pioneering human studies have shown that low-dose ketamine, which acts on brain glutamate receptor pathways (among others), has rapid, robust therapeutic effects in depression and other disorders^8-11^. A randomized controlled clinical trial comparing a single low-dose intravenous ketamine with placebo in OCD patients showed ketamine’s therapeutic effect was rapid (within hours), and half those who received the drug reported remarkable OCD symptom relief (lasting up to 7 days)^12^. A recent rodent study showed that ketamine rapidly reversed a pharmacologically induced perseverative hyperlocomotion behavior^13^. However, the mechanisms underlying ketamine’s therapeutic effect in OCD (and other disorders) are unknown. This knowledge gap is largely due to the fact that no preclinical studies have addressed the circuit effects of ketamine on compulsive behavior in rodent models, a necessary step towards identifying mechanism. Understanding how ketamine modulates neural circuits involved in compulsive behavior will help identify pathways for the rational development of fast-acting therapeutic interventions.

Neuroimaging studies in OCD patients consistently show altered activity of fronto-striatal circuits implicated in action selection, emotion regulation, and cognitive flexibility, although there is conflicting evidence about the directionality of these alterations^14-21^. OCD is associated with dysfunction in two frontal cortical regions: the orbitofrontal cortex (OFC) and dorsal anterior cingulate cortex (dACC). Previous mechanistic work has largely focused on the OFC, a region implicated in assigning value to action outcomes^22^. In contrast, little mechanistic work has addressed the causal role of the dACC in compulsive behavior. The dACC is important for controlling action selection and has also been implicated in fear expression, making it uniquely situated to control both the motor and affective components of OCD^23, 24^. The dACC has been associated with conflict monitoring in OCD^25-27^, and several structural imaging studies have shown decreased ACC volumes in OCD^28, 29^. The fronto-striatal circuit consisting of dACC projections to the dorsal medial striatum (DMS) is of particular interest given its role in controlling cognitive flexibility and goal-oriented behavior, functions that are disrupted in OCD^14, 15, 30^. Alterations in this circuit may lead to perseverative behaviors by disrupting the balance between goal-oriented and habitual behavior^31-33^. Indeed, several studies support a correlation between increased reliance on habitual learning and severity of compulsions^30, 34^. Determining the causal role of the dACC-DMS circuit in compulsive behavior – and how this circuit is modulated by ketamine – could provide critical insights into OCD pathology that are necessary for developing better treatment options.

Rodent studies are beginning to provide insights into cortico-striatal mechanisms of OCD-related behavior^35-37^. One of the best validated rodent models of compulsive behavior is the SAP90/PSD95-associated protein 3 (SAPAP3) knock-out (KO) mouse, which has a prominent compulsive grooming phenotype that is rescued by SRI treatment^38^. Mice lacking SAPAP3 also demonstrate impaired cognitive flexibility, aberrant habit formation, and altered glutamatergic signaling in the dorsal striatum, a region implicated in OCD pathology^38-40^. A recent analysis of post-mortem brains demonstrated reduced expression of the SAPAP3 protein in the striatum of OCD patients, and variants of the *SAPAP3* gene have been associated with early-onset OCD and trichotillomania, another compulsive disorder^41^. Previous studies using this genetic model to investigate fronto-striatal circuits have focused on the role of the OFC, with little work addressing the rodent dACC homolog— the dorsomedial prefrontal cortex (dmPFC)^42, 43^. Given the importance of the dmPFC-DMS circuit for controlling behavioral flexibility, we hypothesized that alterations in the dmPFC-DMS circuit would be associated with compulsive behaviors in this mouse^31^. The SAPAP3 KO mouse thus provides a platform for testing the causal role of the dmPFC-DMS circuit in compulsive behavior and how novel therapeutics such as ketamine modulate compulsive grooming and its underlying neural circuitry.

Here we demonstrate that ketamine rapidly attenuates compulsive grooming behavior in SAPAP3 KO mice. Using *in vivo* fiber photometry recording from dmPFC-DMS projection neurons, we show that SAPAP3 KO mice lack normal modulation of dmPFC-DMS circuit activity associated with grooming behavior in WT mice. Furthermore, ketamine selectively increases activity of this circuit in KO but not WT mice during grooming. Optogenetically mimicking this increased activity by stimulating dmPFC-DMS projections was sufficient to rescue the compulsive grooming phenotype in KO mice. Conversely, optogenetic inhibition of dmPFC-DMS projections increased grooming in WT mice, demonstrating bidirectional control of grooming behavior via this fronto-striatal circuit. These studies demonstrate that ketamine increases activity in a fronto-striatal circuit that causally controls compulsive grooming behavior, suggesting a central role for this circuit in mediating ketamine’s therapeutic effects in OCD.

## Results

### Compulsive grooming is associated with blunted fronto-striatal circuit dynamics

First, we wanted to observe natural neural dynamics in the dmPFC-DMS projection underlying healthy versus compulsive grooming in WT and SAPAP3 KO littermates, respectively. We first confirmed that KO mice had increased grooming compared to WT littermates (**Figure 1a**). To assess circuit activity during behavior, we used fiber photometry to record calcium signals from fronto-striatal projection neurons expressing the calcium indicator GCaMP6m. To selectively express GCaMP6m in dmPFC-DMS projection neurons, we injected a retrograde canine adenovirus carrying Cre recombinase (CAV2-Cre) into the DMS and a Cre-dependent GCaMP6m virus into dmPFC. We then implanted an optical fiber in the dmPFC to record from this DMS-projecting subpopulation (**Figure 1b**). We measured the amplitude (**Figure 1c**) and frequency (**Figure 1d**) of calcium transient peaks during grooming epochs and saw a significant increase in peak frequency in the KO mice. We then generated a z-scored peri-event time histogram (PETH) of dmPFC-DMS projection neuron activity aligned to the onset of grooming behavior. For generating z-scored PETHs we only used grooming epochs that were separated by a minimum of 5 seconds from the preceding grooming epoch to reduce any overlap in signal produced by temporally close grooming bouts (see Methods for details). In WT animals, we observed a decrease in the neural signal with the start of grooming (**Figure 1e**). This dip in activity upon grooming onset was absent in KO mice (**Figure 1f**). To further quantify this genotype difference in neural activity during grooming, we first analyzed the transition into grooming by calculating the area under the curve (AUC) of the PETH immediately before and after grooming onset (in 2-second windows) and saw a significant negative AUC upon grooming in WT animals that was not present in KO animals (**Figure 1g**). We then analyzed neural activity for the entire grooming duration by analyzing the normalized AUC (each grooming epoch divided by the duration of that grooming epoch). We saw that WT animals had a significantly more negative normalized AUC compared to KO animals (**Figure 1h**). To assess whether this circuit is also modulated with cessation of grooming, we then focused on the transition out of grooming behavior. We generated PETHs aligned to the end of grooming epochs and saw that both WT (**Figure 1i**) and KO (**Figure 1j**) animals showed an increase in neural activity immediately after the termination of grooming. We quantified this change by calculating the AUC for 2-second windows immediately before and after the end of a grooming epoch and saw that both WT and KO animals showed a significant increase in neural activity immediately after grooming ended (**Figure 1k**). In summary, WT animals have a sustained decrease in fronto-striatal activity throughout grooming. KO animals, however, selectively lack this robust dip in neural activity at grooming onset, suggesting altered encoding of grooming initiation and/or maintenance in this circuit. Furthermore, both WT and KO animals had a significant increase in neural activity after grooming ends, potentially indicating that different circuits control grooming initiation and termination in the dmPFC. These results together support the hypothesis that dmPFC-DMS circuit modulation is involved in normal grooming behavior, and that engagement of this circuit is markedly altered in compulsive groomers.

**Figure 1.**
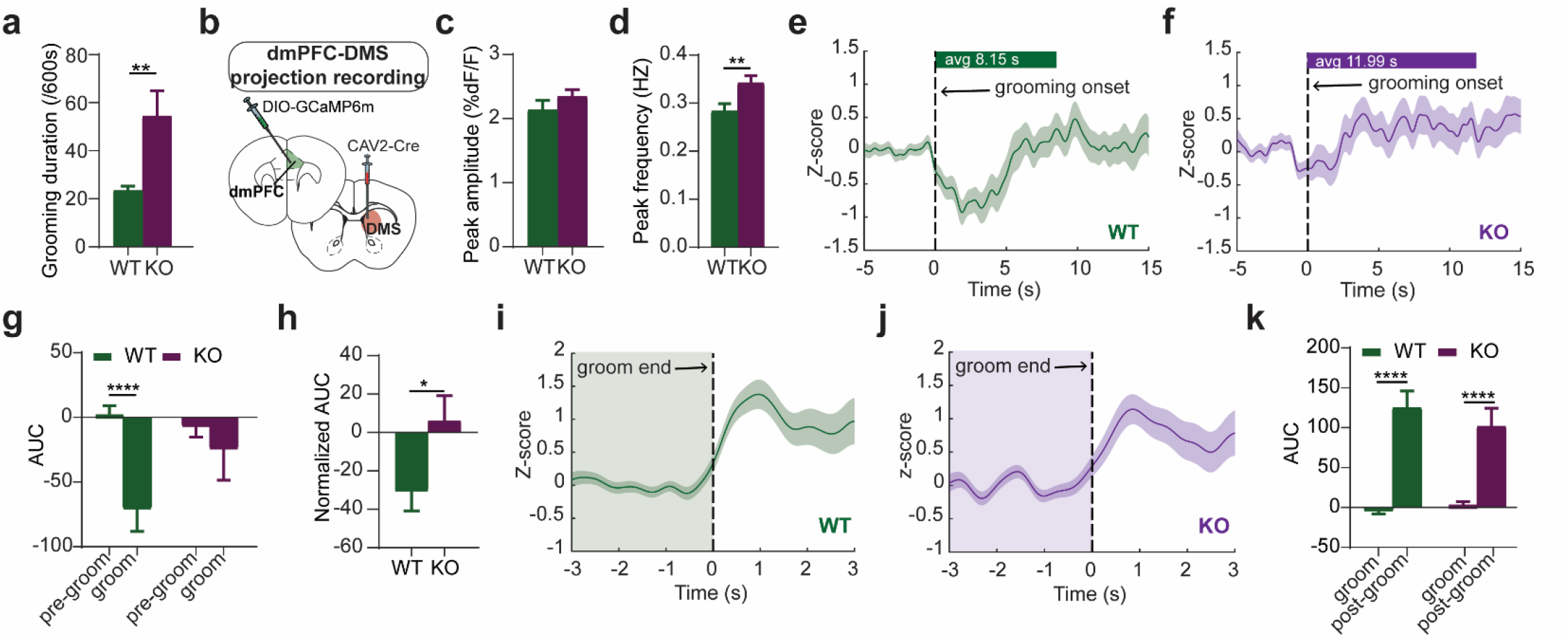
Grooming-associated changes in fronto-striatal activity. **(a)** KO mice groom significantly more than WT littermates (unpaired two-tailed t-test P < 0.01; WT = 15, KO = 15). **(b)** Mice were injected with CAV2-Cre in the DMS and Cre-dependent GCaMP6m in the dmPFC, with optical fibers also implanted in the dmPFC. **(c)** There is no genotype difference in peak amplitude of calcium transients. **(d)** KO mice have a significantly higher frequency of calcium transients than WT mice in dmPFC-DMS projection neurons during grooming (unpaired two-tailed t test P < 0.01; WT = 130 grooming epochs from 11 animals, KO = 118 grooming epochs from 12 animals). **(e)** Peri-event time histogram (PETH) of averaged z-score calcium signals aligned to the start of grooming show that WT mice (green line) have a dip in signal during grooming (green bar above represents average length of grooming epochs for WT). **(f)** KO mice lack this change in z-score during grooming (purple line, purple bar above represents average length of grooming epochs for KO). **(g)** Area under the curve (AUC) analysis of PETHs shows that WT mice have a significant decrease in AUC during grooming while KO mice do not (Two-Way RM ANOVA: interaction P < 0.05, genotype P >0.05, Pre-groom versus groom P <0.01; Sidak’s multiple comparisons: **** = P < 0.0001). **(h)** Normalized AUC analysis shows that the decrease in calcium signal during grooming is significantly different between WT and KO animals (unpaired two-tailed t test P < 0.05; WT = 113 groom epochs from 11 animals, KO = 73 groom epochs from 12 animals). **(i)** PETH of averaged z-score signals aligned to the end of grooming show that WT mice have an increase in calcium signal after grooming ends. **(j)** KO mice have a similar increase in signal after grooming ends. **(k)** AUC analysis shows that both WT and KO mice have a significant increase in neural activity post-grooming (Two-Way RM ANOVA: interaction P > 0.05, genotype P >0.05, groom versus post-groom P <0.0001; Sidak’s multiple comparisons: **** = P < 0.0001).

### Ketamine rescues compulsive grooming behavior

After establishing baseline differences in dmPFC-DMS circuit activity between normal and compulsive grooming, we tested whether ketamine could rescue the behavioral and circuit deficits observed in SAPAP3 KO mice. We first performed a behavioral pharmacology experiment to determine whether ketamine could attenuate compulsive grooming in SAPAP3 KO mice similar to its rapid therapeutic effect observed in human OCD patients. First, we established baseline grooming levels for naïve SAPAP3 KO mice and their WT littermates. We then divided the mice into two experimental groups, to receive either saline or ketamine (30 mg/kg i.p.) injection. To validate that ketamine injections were successful, we measured locomotor activity for 10 minutes post-injection (**Supplemental Figure 1**), as ketamine has a known acute hyperlocomotor effect within ~20 minutes of injection^44^. We then recorded grooming behavior and locomotor activity at four additional time points over the course of a week: 1 hour, 1 day, 3 days, and 7 days post-injection to assess lasting effects of ketamine on compulsive grooming and locomotion beyond the range of its acute psychomotor effects (**Figure 2a** and **Supplemental Figure 1b**, respectively). These time points were chosen to mirror the time points at which a therapeutic effect of ketamine was observed in human OCD patients. We quantified both the number of grooming bouts that mice initiated (grooming frequency) as well as the total time spent grooming (grooming duration). At baseline, SAPAP3 KO mice had significantly higher grooming frequency than WT littermate controls. Ketamine injection markedly attenuated this compulsive grooming phenotype, restoring normal grooming levels that were indistinguishable from WT animals (**Figure 2b**). Specifically, KO mice that received ketamine had significantly reduced grooming frequency at 1 hour and 1 day post-injection that was indistinguishable from WT grooming frequency. This attenuated KO grooming gradually returned back to baseline levels by day 7. Ketamine injection had no effect on grooming frequency in WT littermates, and saline injection had no effect on either group. We then plotted these data as a change from baseline to further demonstrate within-genotype effects of ketamine. We saw significant differences between experimental groups and a significant interaction between group and time post-injection. Post-hoc analysis revealed that KO-saline and KO-ketamine groups significantly differed in grooming frequency at 1 hour and 1 day time points (**Figure 2c**), whereas there was no difference in grooming frequency between WT-saline and WT-ketamine groups at any time point (**Figure 2d**). A similar pattern of results emerged when we quantified total grooming duration (**Figure 2e-g**). These data indicate that ketamine selectively attenuates compulsive over-grooming in KO animals without disrupting normal WT grooming behavior.

**Figure 2.**
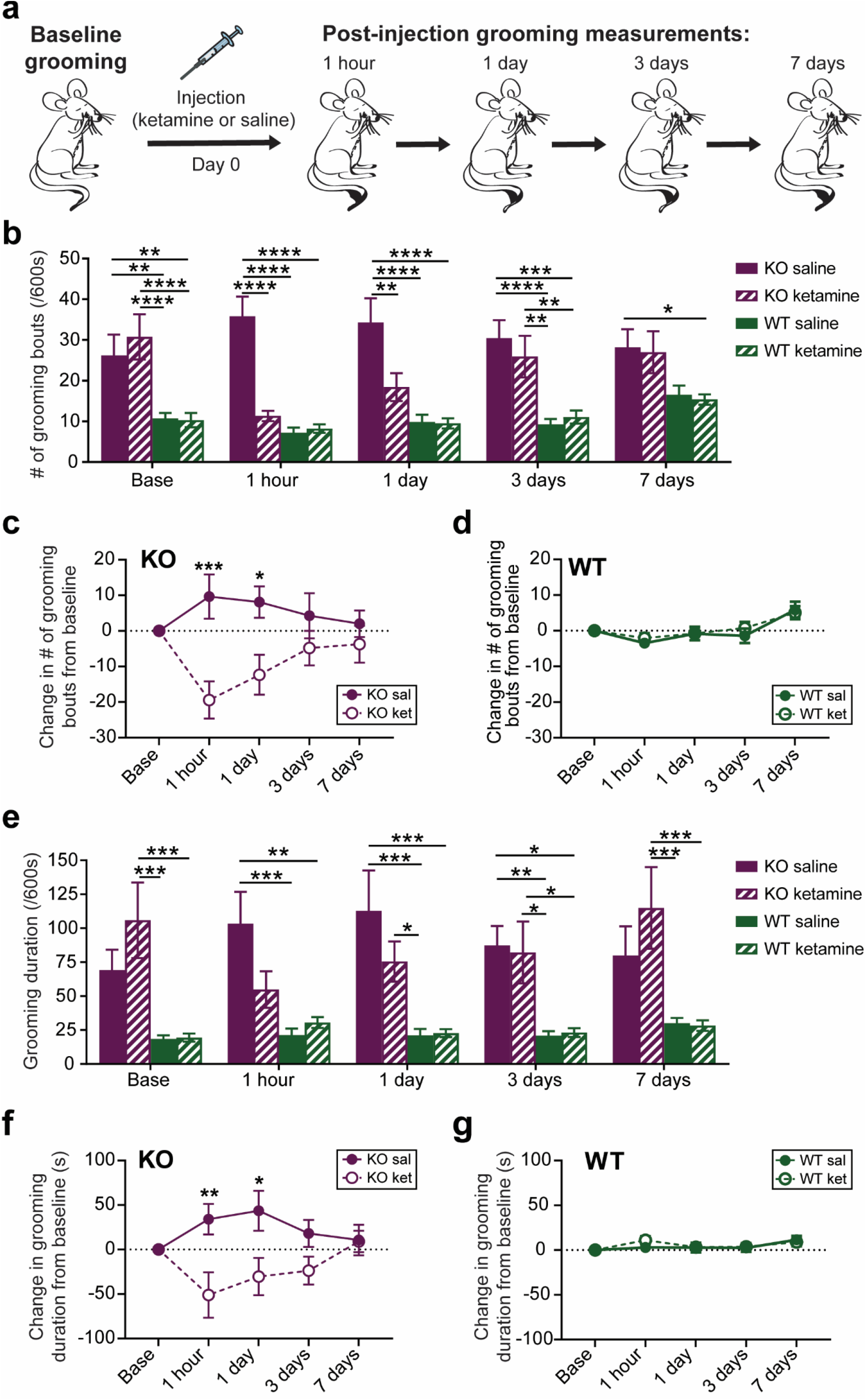
Ketamine reduces compulsive grooming in SAPAP3 KO mice. **(a)** Baseline grooming behavior was quantified in both WT and KO mice. Both genotypes then received either saline or ketamine (30 mg/kg), and grooming measurements were collected over 7 days after injection. **(b)** Both groups of SAPAP3 KO mice (purple) had significantly increased baseline grooming frequency compared to WT controls (green). Ketamine administration significantly reduced grooming frequency in KO animals 1 hour and 1 day post-injection (purple striped bars), abolishing the baseline difference between the KO ketamine group and both WT groups, and producing a significant difference between KO saline and KO ketamine groups (Two-Way RM ANOVA: interaction P < 0.001, time P < 0.05, experimental group P < 0.0001; Tukey’s multiple comparisons: * = P < 0.05, ** = P < 0.01, *** = P < 0.001, **** = P < 0.0001). **(c)** KO mice given ketamine (open circles) show a significant decrease in grooming frequency from baseline compared to KO mice given saline (filled circles; Two-Way RM ANOVA: interaction P<0.01, time P>0.05, experimental group P<0.05; Sidak’s multiple comparisons: * = P < 0.05, *** = P < 0.001). **(d)** WT mice show no significant change in grooming frequency between saline and ketamine groups (Two-Way RM ANOVA: interaction P > 0.05, time P > 0.05, experimental group P > 0.05). **(e)** Similar to grooming frequency, ketamine attenuated grooming duration in KO animals (N = 12) at 1 hour postinjection, abolishing the significant difference in grooming compared to both WT groups at baseline (saline N = 12, ketamine N = 13). Saline injection significantly increased KO grooming (solid purple bars) compared to WT controls 1 hour, 1 day, and 3 days post-injection, returning to baseline levels on day 7 (Two-Way RM ANOVA: interaction P < 0.01, time P > 0.05, experimental group P < 0.0001; Tukey’s multiple comparisons: * = P < 0.05, ** = P < 0.01, *** = P < 0.001, **** = P < 0.0001). **(f)** Changes in grooming behavior from baseline showed that the KO ketamine group had significantly decreased grooming compared to the KO saline group (Two-Way RM ANOVA: interaction P < 0.05, time P > 0.05, experimental group P < 0.01; Sidak’s multiple comparisons: * = P < 0.05, ** = P < 0.01). **(g)** WT mice displayed no significant difference in grooming between treatment groups (Two-Way RM ANOVA: interaction P > 0.05, time P < 0.01, experimental group P > 0.05).

### Ketamine selectively increases in vivo fronto-striatal projection activity in compulsive groomers

Previous studies suggest that ketamine increases glutamatergic transmission and alters excitatory/inhibitory balance within the mPFC^45, 46, 47^. However, our baseline photometry recordings (Figure 1) indicated that KO animals had increased fronto-striatal activity compared to WT animals during grooming, leading us to hypothesize that the therapeutic effect of ketamine might be associated with decreasing activity in this projection to resemble WT activity patterns during grooming. We therefore sought to determine how ketamine specifically affects recruitment of fronto-striatal projection neurons during compulsive behavior. We used *in vivo* fiber photometry to record from dmPFC-DMS projection neurons during grooming, focusing on measuring circuit activity associated with the therapeutic effect of ketamine 24 hours post-injection (**Figure 3a**). To probe these mechanisms, we first injected mice with saline and then 24 hours later analyzed circuit activity during grooming. One week after saline, we injected mice with 30 mg/kg ketamine. This injection order allowed us to assess within-animal circuit changes without concern of potential long-term carryover effects from ketamine exposure that could occur with a Latin squares crossover design, as observed in human studies^12^. We first confirmed the change in grooming induced by ketamine relative to saline injection and saw that KO animals significantly reduced their grooming compared to WT animals post-ketamine (**Figure 3b**). We then quantified the peak frequency and peak amplitude of calcium transients during grooming behavior after saline and ketamine injection. We saw no differences between genotypes or drug conditions in the peak frequency during grooming behavior (**Figure 3c**). However, in KO animals ketamine significantly increased peak amplitude of calcium transients compared to KO-saline, WT-saline, and WT-ketamine groups during grooming epochs, while no change was observed for WT animals between treatment conditions (**Figure 3d**). Next we generated PETHs of dmPFC-DMS projection activity aligned to grooming onset. WT-saline and WT-ketamine groups both showed a dip in neural activity at grooming onset (**Figure 3e**). The KO-saline group appeared to have a subtle dip in neural activity at grooming onset, whereas the KO-ketamine group showed a large increase in neural activity at grooming onset (**Figure 3f**). Quantifying the normalized AUC (as in Figure 1h) confirmed that the KO-ketamine group had significantly increased neural activity during grooming compared to KO-saline, WT-saline, and WT-ketamine groups (**Figure 3g**). We then generated PETHs aligned to the end of grooming epochs and saw that WT-saline and WT-ketamine groups had a similar increase in neural activity immediately after the termination of grooming behavior (**Figure 3h**). KO-saline mice also had an increase in neural activity after grooming ended, but this effect appeared to be disrupted in KO-ketamine mice (**Figure 3i**). To quantify these differences, we calculated the AUC immediately before and after the end of grooming and saw a significant increases in AUC upon grooming end for WT-saline, WT-ketamine, and KO-saline groups, whereas the KO-ketamine group did not show a significant increase in AUC (**Figure 3j**). These results indicate that loss of SAPAP3 sensitizes dmPFC-DMS circuitry to ketamine-induced increases in activity, while this circuitry appears resistant to such effects in WT animals. Supporting the notion that ketamine exerts its behavioral effect via altered *in vivo* recruitment of dmPFC neurons during grooming, *ex vivo* slice electrophysiology recordings showed no effect of genotype or drug treatment on intrinsic excitability of dmPFC neurons (**Supplemental Figure 2**). Additionally, because ketamine did not restore KO dmPFC-DMS projection activity to WT levels, but instead further increased activity in this projection, it is possible that the higher fronto-striatal activity we observed during pre-injection baseline grooming (Figure 1d,f) may represent a compensatory mechanism in KO animals.

**Figure 3.**
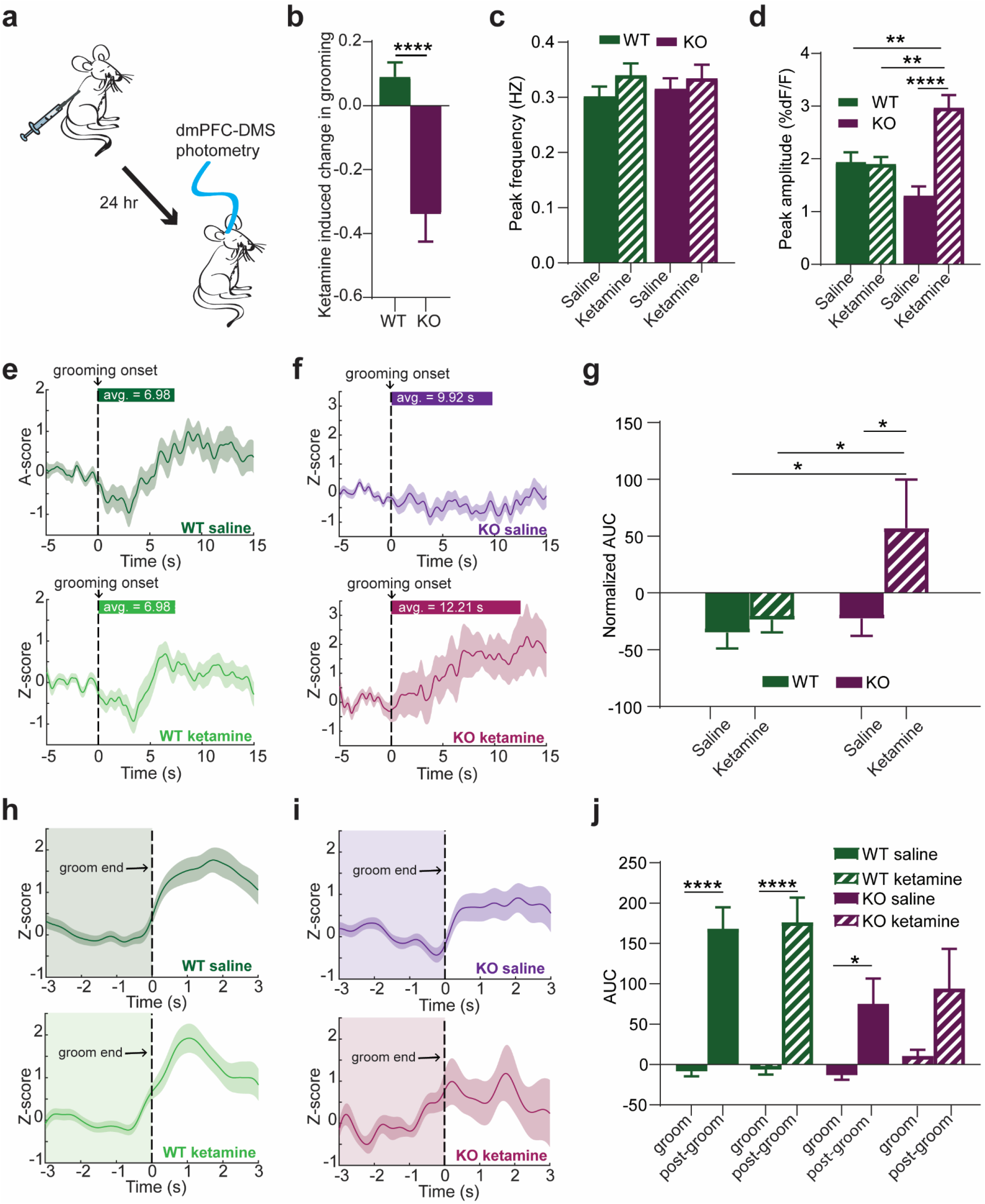
Effect of ketamine on fronto-striatal circuit activity. **(a)** Schematic showing fiber photometry recording of dmPFC-DMS projection neurons 24 hours after mice were injected with saline or ketamine. **(b)** KO mice show a significant decrease in grooming duration compared to WT mice after ketamine injection (unpaired two-tailed t test P<0.0001, WT = 16 animals, KO = 15 animals). **(c)** No change in the frequency of calcium transients between saline and ketamine groups (Two-Way ANOVA; interaction P >0.05, genotype P >0.05, drug P >0.05; WT = 66 saline groom epochs and 70 ketamine groom epochs from 8 animals, KO saline = 69 groom epochs from 12 animals, KO ketamine = 25 groom epochs from 11 animals). **(d)** Ketamine significantly increases the amplitude of calcium transients during grooming epochs in KO animals but not WT animals 24 hours postinjection (Two-Way ANOVA: interaction P < 0.0001, drug P < 0.0001, genotype P > 0.05; Tukey’s multiple comparisons: * = P < 0.05, ** = P < 0.01, *** = P < 0.001, **** = P < 0.0001). **(e)** Peri-event time histograms of averaged z-score signals aligned to the start of grooming show that WT mice have a dip in calcium signal during grooming (green bar above represents average length of grooming epoch for WT) after both saline (top panel) and ketamine injections (bottom panel). **(f)** KO mice show a shallow reduction in neural signal with the onset of grooming after saline injection (top panel) and an increase in signal during grooming after ketamine injection (bottom panel). **(g)** Area under the curve (AUC) quantification for each grooming epoch shows that the KO ketamine group is the only one with a positive AUC that is significantly greater than KO saline, WT saline, and WT ketamine groups (Two-Way ANOVA; interaction P = 0.06, genotype P < 0.05, drug effect P < 0.05; Tukey’s multiple comparison test: * = P < 0.05; WT = 51 grooming epochs with saline and 55 grooming epochs with ketamine from 11 animals, KO saline = 46 grooming epochs from 12 animals, KO ketamine =17 grooming epochs from 11 animals). **(h)** Peri-event time histograms of averaged z-score signals aligned to the end of grooming show that WT saline (top panel) and WT ketamine (bottom panel) groups both have an increase in neural signal immediately after the termination of grooming. **(i)** KO saline animals have an increase in neural signal immediately after the termination of grooming (top panel), but this effect appears to be disrupted in KO ketamine animals (bottom panel). **(j)** AUC analysis confirms a significant increase in neural signal after groom end in WT saline, WT ketamine, and KO saline groups (Two-Way RM ANOVA: interaction P = 0.0505, genotype P > 0.05, groom versus post groom P < 0.0001; Sidak’s multiple comparisons: * = P < 0.05, **** = P < 0.0001).

### Optogenetic inhibition of dmPFC-DMS projections induces over-grooming in WT animals

Our observation that WT animals have a dip in dmPFC-DMS projection activity at the onset of grooming suggested that reduced activity in this circuit may be permissive of grooming. To test this hypothesis, we used optogenetic inhibition to decrease activity in the dmPFC-DMS projection and examine whether this manipulation could causally increase grooming behavior in WT animals. We injected WT mice in the dmPFC with either eYFP as control or halorhodopsin (eNpHR3.0) under control of the CaMKIIα promoter and implanted optical fibers in the DMS to bilaterally inhibit fronto-striatal terminals (**Figure 4a**). Grooming behavior was scored during a 20-minute session consisting of a 5-minute baseline period, followed by 10 minutes with the laser on, and an additional 5 minutes after the laser was turned off. When the laser was turned on to inhibit the dmPFC-DMS projection, grooming behavior increased in the eNpHR3.0 group, whereas eYFP mice maintained constant levels of grooming across the entire experimental time window (**Figure 4b**). The eNpHR3.0 group showed a shorter latency to their first grooming bout after the laser was turned on (**Figure 4c**), but still had a delay of ~30 seconds to grooming onset. The fact that this increase in grooming behavior was not time-locked to the onset of the laser suggests that inhibiting this projection is permissive of grooming rather than directly driving the behavior. We averaged the amount of time spent grooming across 5-minute epochs of laser on versus laser off and saw that in the eNpHR3.0 group grooming significantly increased for the first 5 minutes while the laser was on (**Figure 4d**). Analysis revealed a significant main effect of virus with a significant interaction between virus and laser. At baseline, there were no grooming differences between eYFP and eNpHR3.0 groups. Mice expressing eNpHR3.0 but not eYFP had a significant laser-evoked increase in grooming duration from baseline (**Figure 4e**). These results indicate that inhibiting the dmPFC-DMS projection produces a hyper-grooming phenotype in WT mice.

**Figure 4.**
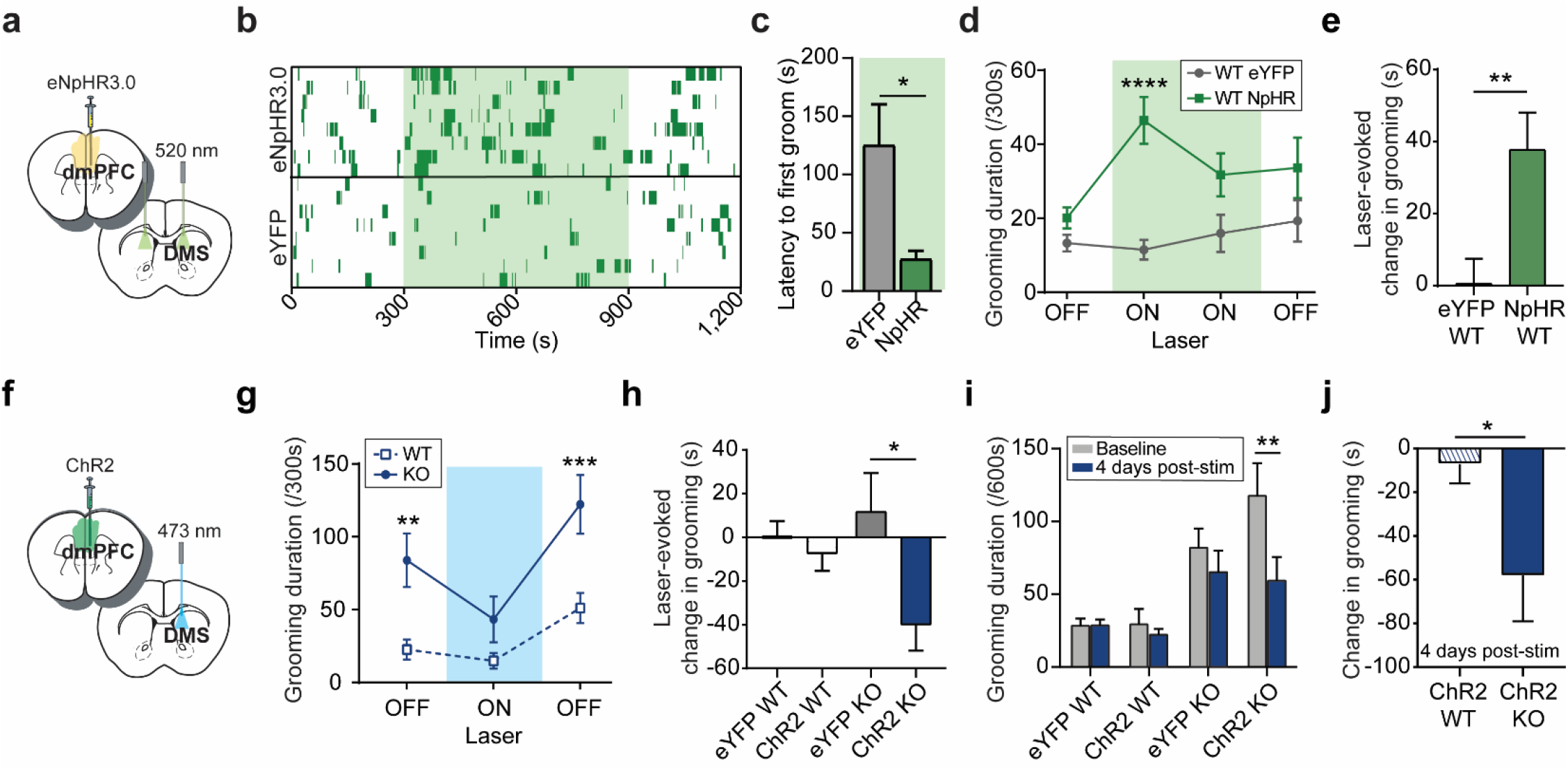
Bidirectional control of compulsive behavior via optogenetic manipulation of dmPFC-DMS projections. **(a)** WT mice were bilaterally injected with CaMKIIα-eNpHR3.0 virus in the dmPFC and implanted bilaterally with optical fibers in the DMS. **(b)** Raster-like plot of grooming behavior for individual animals across experimental timeline showing that dmPFC-DMS projection inhibition increases grooming behavior (top: eNpHR3.0; bottom: eYFP control). Each row represents one animal. Dark green dashes indicate grooming timestamps and light green shading indicates when laser was on. **(c)** Mice expressing eNpHR3.0 began grooming significantly sooner than mice expressing eYFP after laser onset (unpaired two-tailed t-test P < 0.05). **(d)** Mice expressing eNpHR3.0 (N = 8) showed increased grooming during laser on epochs that was significantly higher than eYFP controls (N = 8), with a significant main effect of virus, a significant interaction between laser and virus, and a trend in the main effect of laser. Multiple comparisons revealed a significant difference between eYFP and eNpHR3.0 groups during the first 5-minute laser on block (Two-way RM ANOVA: interaction P < 0.05, laser P = 0.0841, virus group P < 0.001; Sidak’s multiple comparisons: **** = P < 0.0001). **(e)** Bar graph quantifying data from (d) showing that eNpHR3.0 mice have a significant laser-evoked increase in grooming compared to eYFP mice (unpaired two-tailed t test P < 0.01). **(f)** KO and WT mice were unilaterally injected with CaMKIIα-ChR2 virus in the dmPFC and implanted with an optical fiber in the DMS. **(g)** At baseline, SAPAP3 KO mice expressing ChR2 (N = 15) have significantly higher levels of grooming than WT mice expressing ChR2 (N = 18). This difference is abolished during laser stimulation (blue shaded bar), and returns to significantly higher levels again once the laser is turned off (Two-way RM ANOVA: interaction P < 0.05, laser P < 0.0001, virus group P < 0.01; Sidak’s multiple comparisons: ** = P < 0.01, *** = P < 0.001). **(h)** Bar graph quantifying data from (g) showing that KO ChR2 mice (N = 15) have a significant reduction in grooming behavior when the laser is on compared to KO eYFP mice (N = 10) (unpaired two-tailed t test P < 0.05). There is no significant difference in grooming behavior between WT eYFP (N = 11) and WT ChR2 (N = 18) groups (unpaired two-tailed t test P > 0.05). **(i)** Following two weeks of repeated stimulation, KO ChR2 mice had a significant and sustained reduction in grooming behavior four days after the last laser stimulation session (Two-way RM ANOVA: interaction P < 0.05, time P < 0.0001, experimental group P < 0.01; Sidak’s multiple comparisons: ** = P < 0.01; WT eYFP N = 11, WT ChR2 N = 13, KO eYFP = 10, KO ChR2 = 10). **(j)** Four days after the final stimulation after two weeks of repeated stimulation, KO ChR2 mice show a significantly greater decrease from baseline grooming behavior than WT ChR2 mice (unpaired two-tailed t test P < 0.05).

### Stimulation of dmPFC-DMS projections rescues compulsive grooming in SAPAP3 KO animals

Given that the therapeutic effect of ketamine was associated with increased neural activity in fronto-striatal projection neurons during grooming in KO animals, we reasoned that optogenetically elevating activity in this circuit might likewise have a therapeutic effect on compulsive grooming. We injected channelrhodopsin-2 (ChR2) under control of the CaMKIIα promoter into the dmPFC and implanted an optical fiber in the DMS (**Figure 4f**). We recorded grooming behavior for a 5-minute baseline period, a 5-minute laser on period (10 Hz stimulation), and a final 5-minute laser off period. At baseline, we saw a significant difference in grooming levels between WT-ChR2 and KO-ChR2 animals. When the laser was on, KO-ChR2 grooming behavior was reduced to WT levels. Upon termination of the laser, KO-ChR2 grooming returned to significantly higher levels than the WT-ChR2 mice, demonstrating that stimulation of dmPFC terminals in the DMS had an acute therapeutic effect on aberrant KO grooming behavior (**Figure 4g**). Interestingly, stimulation of dmPFC terminals did not alter the number of grooming bouts that KO mice engaged in, demonstrating that stimulation of dmPFC only altered duration of grooming (**Supplemental Figure 3**). Laser stimulation had no effect on grooming of KO-eYFP controls or WT-ChR2 and WT-eYFP groups (**Figure 4h**). Since stimulating this projection had an acute effect of attenuating compulsive grooming, we tested whether repeated stimulation of this projection could produce a lasting therapeutic effect. Mice received three additional stimulation sessions over two weeks, totaling 45 minutes of laser stimulation. We then scored grooming behavior in the absence of laser four days after the last stimulation to look for lasting therapeutic effects. KO-ChR2 cumulative grooming behavior was indeed significantly reduced after this repeated stimulation compared to pre-stimulation baseline grooming behavior, an effect that was not present in KO-eYFP, WT-eYFP, and WT-ChR2 groups (**Figure 4i**). Specifically, when we compared the change in grooming duration after repeated stimulation between KO-ChR2 and WT-ChR2 groups, we saw that KO-ChR2 had a significantly greater reduction in grooming behavior relative to WT-ChR2 (**Figure 4j**). These optogenetic experiments together demonstrate causal control of compulsive grooming behavior via manipulation of the dmPFC-DMS projection. Furthermore, these results indicate that minimal repeated stimulation of this projection may have lasting therapeutic effects on compulsive behavior.

## Discussion

Cortico-striatal circuits are heavily implicated in the pathophysiology of OCD, although few previous mechanistic studies in rodents have interrogated the role of the dmPFC-DMS circuit in compulsive behavior, despite its known role in OCD-relevant processes such as cognitive flexibility and action-outcome learning. Here we identify how activity of the dmPFC-DMS circuit is altered in SAPAP3 KO mice that display compulsive grooming behavior. Furthermore, we describe how ketamine affects compulsive grooming and its underlying dmPFC-DMS circuit modulation. To our knowledge, this is the first study to use ketamine as a tool to probe fronto-striatal circuit mechanisms underlying compulsive grooming, and one of only two studies to demonstrate ketamine’s therapeutic effect in rodent models of compulsive behavior (the other study recently demonstrated ketamine reversal of a pharmacologically induced perseverative hyperlocomotion)^13^. We show that ketamine provides fast-acting therapeutic relief in SAPAP3 KO mice, reducing both the frequency of grooming bouts and cumulative time spent grooming. This therapeutic effect remained significant 24 hours after ketamine injection and then gradually tapered off over the course of 7 days post-injection. The time course of this anti-compulsive effect of ketamine in SAPAP3 KO mice was strikingly similar to that observed in clinical studies of ketamine in OCD, which found immediate symptom relief that lasted up to 7 days^12^. Our rodent results thus corroborate human studies identifying ketamine as a promising candidate for a fast-acting pharmacological treatment for OCD. These experiments establish the SAPAP3 KO mice as a preclinical translational model for testing ketamine metabolites and related pharmacologic agents to screen for drugs that may have a similar therapeutic profile for OCD-like behavior with fewer side effects^48, 49^.

To determine circuit changes associated with the anti-compulsive effect of ketamine, we used *in vivo* fiber photometry recording of fronto-striatal projection neurons to assess genotype- and drug-related differences in dmPFC-DMS circuit activity during grooming. Our baseline fiber photometry recordings showed several key differences in fronto-striatal circuit activity between WT and SAPAP3 KO mice. WT fronto-striatal projection neurons had a sharp drop in activity at the onset of grooming, but no such modulation was observed in KO mice. The maintained neural activity during grooming observed in KO mice appeared to be driven by an increased number of calcium transients. Interestingly, upon transition out of grooming, both WT and KO mice showed an increase in neural activity. For WT animals this increase in neural activity immediately after grooming is not surprising considering the sustained decrease in activity observed during grooming. However, the increased signal upon termination of grooming in KO animals was unexpected given the lack of decreased neural activity at grooming onset. This could indicate that, with respect to grooming behavior, dmPFC-DMS projection neurons have two states (low activity permits grooming whereas high activity suppresses grooming). Loss of the low activity state with maintenance of the high activity state in KO mice could represent compensatory mechanisms in the dmPFC to try to modulate compulsive grooming behavior. Furthermore, these two activity states could be driven by different synaptic inputs onto the dmPFC, which may be differentially dysregulated in KO mice. For example, in another mouse model of compulsivity, repetitive checking behavior is driven by reduced mPFC glutamatergic activity due to feed-forward inhibition from BLA inputs^50^. In contrast to these *in vivo* circuit differences, *ex vivo* prefrontal slice recordings did not reveal any genotype differences in basal cellular properties of dmPFC neurons. These results suggest that ketamine likely exerts its behavioral effect on grooming not via altered intrinsic excitability of dmPFC cells, but rather via altered *in vivo* recruitment of dmPFC neurons during compulsive behavior. Furthermore, these slice results support the hypothesis that the *in vivo* activity differences we observed between WT and KO mice may be the result of altered presynaptic inputs to dmPFC or altered postsynaptic release in the striatum. For example, other studies have demonstrated altered endocannabinoid signaling in the striatum of SAPAP3 KO mice, which could affect local release properties of glutamatergic projections without changing overall prefrontal cell body activity^51^.

Based on our baseline neural recordings, we originally hypothesized that ketamine administration in KO animals might restore the dip in fronto-striatal activity observed in WT animals during grooming. Instead, ketamine significantly increased dmPFC-DMS projection activity in KO but not WT animals. This suggests that the lack of a grooming-related dip in neural activity during baseline recordings in KO mice may represent compensatory activity in this projection to counteract hyperactive downstream motor circuitry (e.g., DLS or central striatum) that directly drives compulsive grooming^52^, while the ketamine-induced increase in glutamatergic transmission may allow for dmPFC-DMS projection neurons to more successfully restore normal behavior. Furthermore, it seems unlikely that the WT dip in fronto-striatal activity is directly driving grooming behavior but may instead represent a permissive signal for grooming. The timing of our optogenetic inhibition effect corroborates this idea: inhibiting the dmPFC-DMS projection in WT mice did not immediately drive grooming behavior when the laser was turned on, but rather gradually increased the probability of grooming episodes, consistent with a permissive role of this circuit. Additionally, saline injection, which trended toward exacerbating grooming duration in KO animals, reduced KO fronto-striatal activity during grooming, further supporting a model in which decreased fronto-striatal projection activity promotes the initiation of grooming behavior. Therefore the dmPFC-DMS projection neurons may represent a permissive signal that normally promotes grooming behavior through decreased activity, but is disrupted in KO mice. A recent study demonstrated that cortico-striatal limbic loops, such as the dmPFC-DMS projection, are capable of modulating cortico-striatal motor loops, providing a potential anatomical substrate for permissive dmPFC-DMS gating of DLS that can be causally tested in future studies^53^. The additional boost in prefrontal glutamatergic signaling provided by ketamine may therefore further dampen the dmPFC’s permissive signal to a threshold that is sufficient to overcome dysregulated downstream grooming motor circuitry and restore healthy grooming levels^54^. Indeed, both the pharmacological (ketamine) and optogenetic manipulations we employed that increase activity in the dmPFC-DMS projection had a convergent effect of decreasing compulsive grooming in KO mice (**Figure 5**), while manipulations that reduced dmPFC-DMS activity increased grooming behavior. Interestingly, others have demonstrated that reductions in neural activity of dmPFC projections to the brainstem could induce compulsive alcohol drinking behavior in mice, supporting the idea that disengagement of the dmPFC contributes to compulsive action^55^.

**Figure 5.**
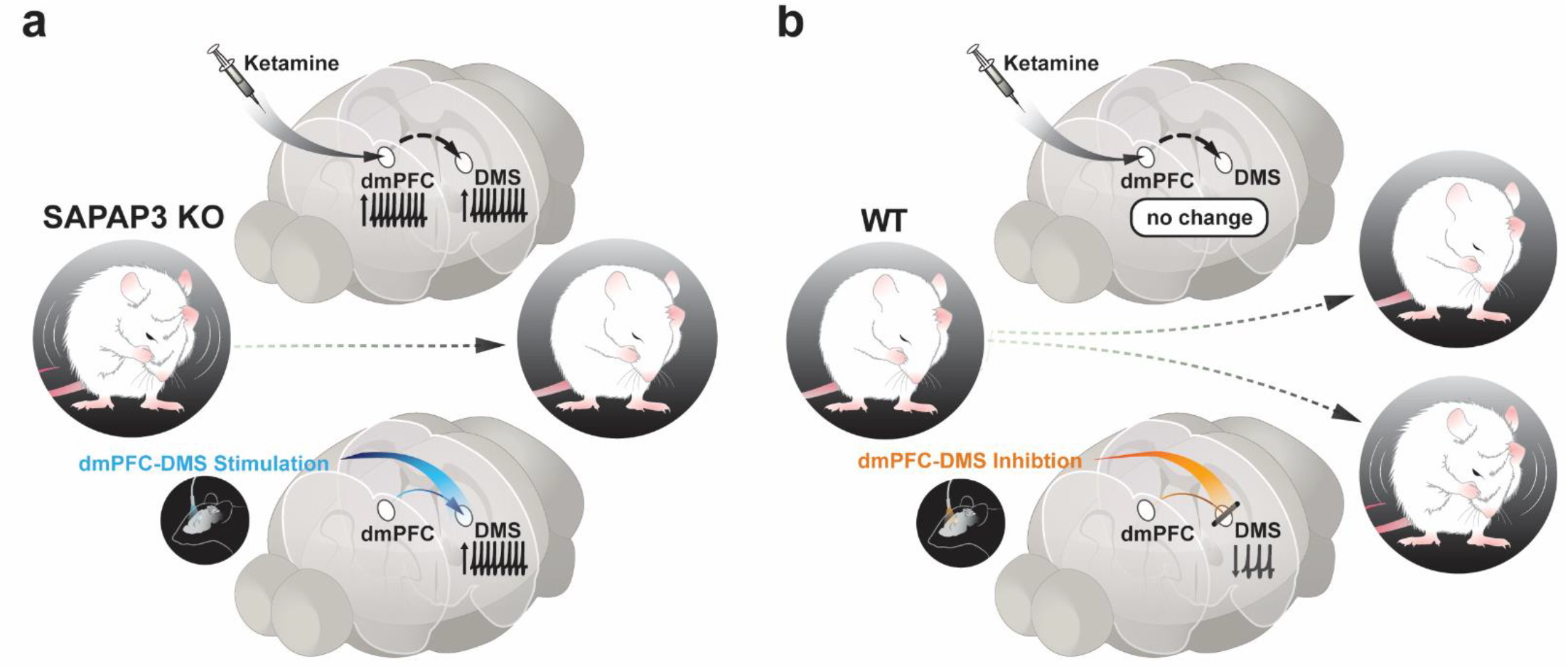
Increasing dmPFC-DMS projection neuron activity via ketamine or optogenetic stimulation rescues compulsive grooming behavior. Graphical summary of the paper showing: **(a)** Ketamine produces a decrease in grooming behavior that is correlated with an increase in dmPFC-DMS projection neuron activity in SAPAP3 KO mice. This rescue of compulsive grooming behavior is causally reproduced by selectively stimulating dmPFC-DMS projection neurons. **(b)** Ketamine has no effect on grooming behavior or dmPFC-DMS circuit activity in WT mice, though optogenetic inhibition of dmPFC-DMS projections in WT mice was sufficient to induce increased grooming.

Several rodent studies have highlighted the importance of the orbitofrontal cortex, ventral striatum, and amygdalo-striatal projection in compulsive behavior^35, 50^. We now add to this growing body of scientific work by demonstrating the causal role of the dmPFC-DMS circuit in compulsive behavior. Furthermore, we provide the first steps in understanding how ketamine can interact with circuits altered in OCD to restore normal goal-directed behavior, providing a preclinical platform for testing neural circuit mechanisms underlying novel fast-acting treatments for OCD.

## Materials and Methods

### Animals

All experiments were performed under a protocol approved by the Institutional Animal Care and Use Committees at the University of California, San Francisco. Homozygous SAPAP3 knockout (KO) mice and their wild-type (WT) littermates were generated from breeding two heterozygotes that were maintained on a C57BL/6J background. For a subset of optogenetic experiments, wild-type C57BL/6J mice purchased from The Jackson Laboratory were also used. The over-grooming phenotype in SAPAP3 KOs emerges around 4 months of age, so all mice were at least 120 days old. Both male and female mice were used. All mice were raised in normal light conditions (12:12 light/dark cycle), with *ad libitum* access to food and water. Experiments were performed during the inactive cycle.

### Surgery

Mice were anesthetized using 5% isoflurane at an oxygen flow rate of 1L/min and placed in a stereotaxic apparatus (Kopf Instruments) on top of a heating pad. Anesthesia was maintained with 1.5-2.0% isoflurane for the duration of surgery. Respiration and toe pinch response were monitored closely. Slow-release buprenorphine (0.5mg/kg) and ketoprofen (1.6mg/kg) were administered subcutaneously at the start of surgery. The incision area was shaved and cleaned with ethanol and betadine. Lidocaine (0.5%) was administered topically. An incision was made along the midline, and bregma and lambda were measured to level the skull. Virus was injected and fiber optic ferrule(s) implanted as described below. After securing the ferrule with dental cement, the skin was sutured around the implant and mice recovered in a clean cage atop a heating pad. The following day, mice were monitored for healthy recovery and administered a subsequent dose of ketoprofen.

#### Inhibition of dmPFC-DMS projections neurons

To optogenetically inhibit projections from dmPFC to DMS, 500 nL of AAV5-CaMKIIα-eNpHR3.0-eYFP or AAV5-CaMKIIα-eYFP control virus (UNC Vector Core) was bilaterally injected into the mPFC (+/-0.35 M/L; 1.8 A/P; −2.6 D/V, in mm from bregma) using a 10 μl nanofil syringe (World Precision Instruments) with a 33 gauge beveled needle at a rate of 0.1 μl / min. After waiting 15 minutes, the needle was slowly withdrawn. Two optical fibers in 1.25 mm ceramic ferrules (0.39 NA, 200 μm diameter, Thorlabs) were then implanted bilaterally in the DMS (+/-1.2 M/L; 0.8 A/P; −3 D/V). Fibers were lowered slowly into the DMS and ferrules were secured to the skull using dental cement (C&B Metabond). These surgeries were done on C57BL/6J WT mice (The Jackson Laboratory).

#### Stimulation of dmPFC-DMS projections neurons

For stimulation of projections from dmPFC to DMS, 500 nL of AAV5-CaMKIIα-ChR2-eYFP (Addgene) or AAV5-CaMKIIα-eYFP (UNC Vector Core) (diluted 1:3 in saline) was injected unilaterally into the left dmPFC and an optical fiber was implanted unilaterally in the left DMS, using the same coordinates and ferrules as described above. These surgeries were done on SAPAP3 KO animals and WT littermates as controls.

#### Fiber Photometry

For recording from dmPFC neurons that project to the DMS, 700 nL of a 50/50 mixture of AAV1.hSyn.mCherry (to visualize injection location; UNC Vector Core) and CAV2-Cre (Plateforme de Vectorologie de Montpellier) was injected unilaterally into the DMS (+/-1.4 M/L; 0.8 A/P; −3.5 D/V). To allow CAV2-Cre-induced GCaMP expression in the dmPFC, 1500 nL of AAV1.Syn.Flex.GCaMP6m.WPRE.SV40 (Addgene) was unilaterally injected in the dmPFC (+/-0.35 M/L; 1.9 A/P; −2.5 D/V). An optical fiber stub in a 2.5 mm ceramic ferrule (0.48 NA, 400 μm diameter, Doric Lenses) was then implanted into the dmPFC (+/-0.35 M/L; 1.9 A/P; −2.3 D/V) and secured as described above. These surgeries were done on SAPAP3 KOs and WT littermates.

### Viruses

pAAV.Syn.Flex.GCaMP6m.WPRE.SV40 was a gift from The Genetically Encoded Neuronal Indicator and Effector Project (GENIE) & Douglas Kim (Addgene viral prep # 100838-AAV1; http://n2t.net/addgene:100838; RRID:Addgene_100838). pENN.AAV.CamKII.GCaMP6f.WPRE.SV40 was a gift from James M. Wilson (Addgene viral prep # 100834-AAV5; http://n2t.net/addgene:100834; RRID:Addgene_100834). pAAV-CaMKIIa-hChR2(H134R)-EYFP was a gift from Karl Deisseroth (Addgene viral prep # 26969-AAV5; http://n2t.net/addgene:26969; RRID:Addgene_26969). AAV5-CaMKIIa-eNpHR3.0-eYFP was a gift from Karl Deisseroth and packaged by the UNC Vector Core. AAV5-CaMKIIa-eYFP was a gift from Karl Deisseroth and packaged by the UNC Vector Core. AAV1-hSyn.mCherry was a gift from Karl Deisseroth and packaged by the UNC Vector Core.

### Behavioral Assays

#### Grooming

To observe grooming behavior, mice were placed in a clear plastic cylinder (36 cm diameter) and recorded with two video cameras to have both top and side views. All behavioral assays were done in an open top sound dampened chamber. A blind observer manually scored and timestamped grooming behavior. Locomotor activity (distance traveled, velocity) was recorded and quantified by Ethovision XT software (Noldus). A TTL pulse triggered by the Ethovision XT software was used to synchronize fiber photometry recording data with behavioral data.

#### Behavioral Pharmacology

Grooming behavior for SAPAP3 KOs and WTs was recorded for 10-minute sessions as described above. Baseline grooming behavior was measured, and then mice were split into two groups to receive either saline (0.9% sterile sodium chloride) or ketamine (30 mg/kg, Sigma). The next day mice were injected intraperitoneally with either saline or ketamine and immediately placed in the behavioral rig. Locomotor recordings were taken for 10 minutes post-injection to verify successful ketamine injection through induction of a short-term increase in locomotor behavior. No grooming measurements were taken at this time. Grooming was scored at additional behavioral time points of 1 hour, 1 day, 3 days, and 7 days post-injection.

#### Optogenetics

Baseline measurements were recorded while mice were not connected to a patch cord and were observed moving freely. On experimental days, a patch cord connected to either a green or blue laser was attached to the ferrule on an individual mouse using a ceramic mating sleeve (Thorlabs). For bilateral inhibition of dmPFC-DMS terminals, each trial began with a 5-minute baseline period followed by 10 minutes of laser on and 5 minutes of laser off post-stimulation. To inhibit projections from dmPFC to DMS using eNpHR3.0, green light was generated by a 532nm laser (Shanghai Laser & Optics Century Co. LTD) (5 mW continuous light on each side). For stimulation of dmPFC-DMS terminals, each session was divided into three epochs: a 5-minute baseline period followed by 5 minutes of laser on and 5 minutes of laser off post-stimulation. To stimulate projections from dmPFC to DMS using ChR2, blue light was generated by a 473 nm laser (Shanghai Laser & Optics Century Co. LTD) (0.5-0.8 mW, 10 Hz, 5 ms pulse width). For stimulation experiments, a subset of animals received three additional days of stimulation totaling 45 minutes spread across two weeks. We then assessed the long-term effects of the laser on grooming behavior four days after the final stimulation when mice were untethered. For all optogenetic experiments a Master-8 (A.M.P.I.) pulse generator controlled by TTL pulses from Ethovision (Noldus) software was used to drive the laser. Laser output was delivered to the animal via an optical fiber (0.39 NA, 200 μm, Thorlabs) connected to a 1×1 fiber optic rotary joint (Doric Lenses) followed by another optical fiber (0.37 NA, 200 μm, Doric Lenses) which was coupled to the cannula through a ceramic sleeve (Thorlabs).

#### Fiber Photometry

Prior to surgery we quantified grooming behavior over 10 minutes for each animal to confirm the increased grooming phenotype in the SAPAP3 KO mice. For dmPFC-DMS fiber photometry recordings, SAPAP3 KOs and WT littermates underwent two days of recording baseline grooming behavior. Each recording session was 20 minutes. Following the end of recording on the second day of baseline, mice received an injection of saline solution (10 mg/mL, i.p.) and were returned to their home cage. Behavioral measurements and fiber photometry recordings then occurred 24 hours later. After 7 days post-saline injection mice were then administered a dose of ketamine (30 mg/kg, i.p.) and returned to their home cage. Behavioral measurements and fiber photometry recordings were taken 24 hours after ketamine injection. Fiber photometry signals were demodulated and analyzed using custom MATLAB code. We restricted the analysis of grooming to epochs longer than 3 seconds. Prior to analysis of the photometry signals post-injection, we determined which KO mice were responders versus non-responders to ketamine by calculating a grooming index to assess change in grooming behavior induced by ketamine ((grooming duration post-ketamine – grooming duration post-saline)/(grooming duration post-ketamine + grooming duration post-saline). We then generated the average and standard deviation for WT animals. KO animals had to have a reduction in grooming behavior post-ketamine was at least 1 standard deviation different from the change in grooming behavior of WT animals to be included the fiber photometry analysis. Only 2 out of 15 KO mice did not meet this criterion. We analyzed the number of calcium transients and the amplitude of these transients that occurred during grooming. We also generated perievent time histograms (PETH) of the neural activity aligned to the beginning and end of grooming bouts and generated area under the curve analysis (AUC) for each grooming epoch. In order for grooming bouts to be included in the groom start PETH analysis, we implemented a 5 second threshold in which the start of the grooming epoch had to be at least 5 seconds after the end of the prior grooming epoch. This allowed us a 5 second window prior to groom start that we knew did not have any signal interference from other grooming epochs that we used to generate the necessary baseline mean signal used to convert the photometry signals into z-score. Additionally we generated a similar 3 second threshold when analyzing the groom end transition in which the start of the next grooming bout had to be greater than 3 seconds after the end of a grooming bout. For groom end we generated the z-scores by using the 3 seconds prior to groom end to generate our baseline signal value.

*In vivo* calcium imaging data were acquired using a custom-built rig based on a previously described setup from the Deisseroth lab (Lerner et al, 2015). This setup was controlled by an RZ5P fiber photometry processor (TDT) and Synapse software (TDT). The RZ5P/Synapse software controlled a 4-channel LED Driver (DC4100, Thorlabs), which in turn controlled two fiber-coupled LEDs: 470 nm for GCaMP stimulation and 405nm to control for artifactual fluorescence (M470F3, M405FP1, Thorlabs). These LEDs were sinusoidally modulated at 210 Hz (470 nm) and 320 Hz (405 nm) and connected to a Fluorescence Mini Cube with 4 ports (Doric Lenses), and the combined LEF output was connected through a fiber optic patch cord (0.48 NA, 400 um, Doric Lenses) to the cannula via a ceramic sleeve (Thorlabs). The emitted light was focused onto a Visible Femtowatt Photoreceiver Module (Model 2151, Newport, DC low) and sampled at 60 Hz. Noldus behavioral analysis was synchronized to the photometry setup using TTL pulses generated every 10 seconds following the start of the Noldus trial.

### Perfusions

Following the conclusion of our behavioral experiments, animals were anesthetized using 5% isoflurane and given a lethal dose (1.0 ml) cocktail of ketamine/xylazine (10 mg/ml ketamine, 1 mg/ml xylazine). They were then transcardially perfused with 10 mL of 1X PBS followed by 10 mL 4% paraformaldehyde (PFA). Brains were left in 4% PFA overnight and then transferred to a 30% sucrose solution until slicing. The brains were frozen and sliced on a sliding microtome (Leica Biosystems) and placed in cryoprotectant in a well-plate. Slices were then washed in 1X PBS, mounted on slides (Fisherbrand Superfrost Plus) and air dried (covered). ProLong Gold antifade reagent (Invitrogen, ThermoFisher Scientific) was pipetted on top of the slices, a cover slip (Slip-rite, ThermoFisher) was placed on top, and the slides were left to dry overnight (covered). Viral injection, fiber photometry cannula implant, and optogenetic cannula implant placements were histologically verified on a fluorescence microscope (Leitz DMRB, Leica). Neural and behavioral data of mice with incorrectly targeted fiber photometry implants were removed from analysis.

### Statistical Analysis

Statistical analyses was performed using GraphPad Prism 8 software package. Statistical significance was set at P < 0.05 for all experimental results. The type of statistical analysis used was determined independently for each experiment and is listed in the relevant figure caption along with the sample number.

## Acknowledgements

We would like to thank Julia Kuhl for her exceptional graphic design work in generating the graphical summary for Figure 5.

## Author Contributions

GLD, CIR, and LAG all contributed to the intellectual conceptualization and design of the overall study. GLD designed, performed, and analyzed data for the behavioral pharmacology and fiber photometry experiments and helped analyze the optogenetic experiments. ARM performed and analyzed optogenetic experiments. AL performed and analyzed slice electrophysiology experiments. GLD and LAG wrote the manuscript and made the figures.

