## Supplemental Material for "Ketamine increases activity of a fronto-striatal projection that regulates compulsive behavior"

### Supplemental Data and Methods

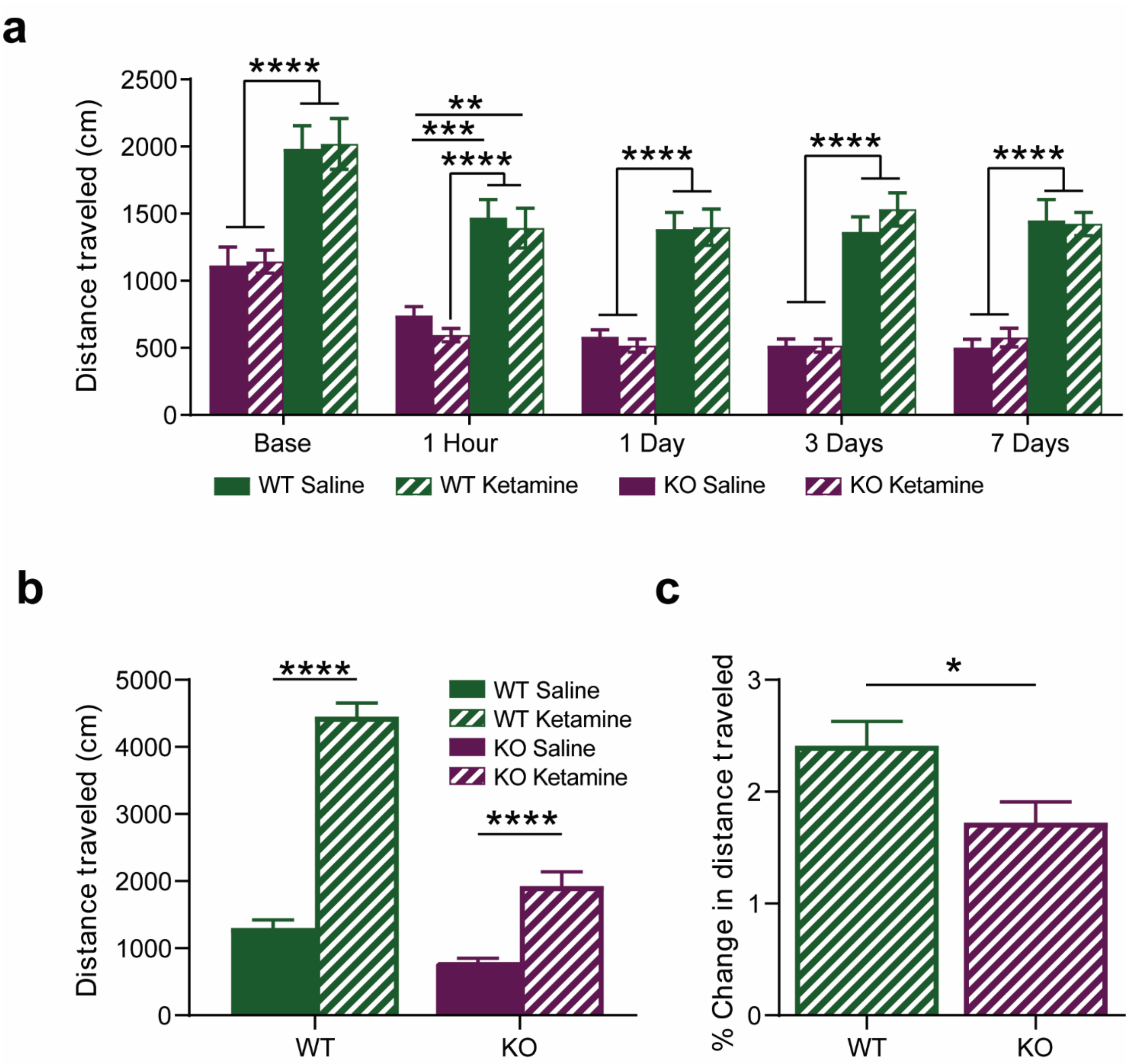


**Supplemental Figure 1. Locomotor effects of ketamine on SAPAP3 KO and WT mice. (a)** Locomotor activity was recorded across the same experimental time window as grooming behavior in Figure 1. KO mice, regardless of treatment, move significantly less than WT littermates (Two-way RM ANOVA: interaction $P > 0.05$, day $P < 0.0001$, experimental group $P < 0.0001$; Tukey's multiple comparisons: ** = $P < 0.01$, *** = $P < 0.001$, **** = $P < 0.0001$). **(b)** Both WT and KO mice treated with ketamine show a significant acute increase in locomotor activity 10 minutes post-injection (Two-way ANOVA: interaction $P < 0.0001$, genotype $P < 0.0001$, experimental group $P < 0.0001$; Sidak's multiple comparisons: **** = $P < 0.0001$). **(c)** However, ketamine induces a larger increase in locomotor behavior (over 200%) in WTs compared to KOs, when ketamine-induced locomotor behavior is normalized to baseline locomotor behavior measures (unpaired two-tailed t-test $P < 0.05$).

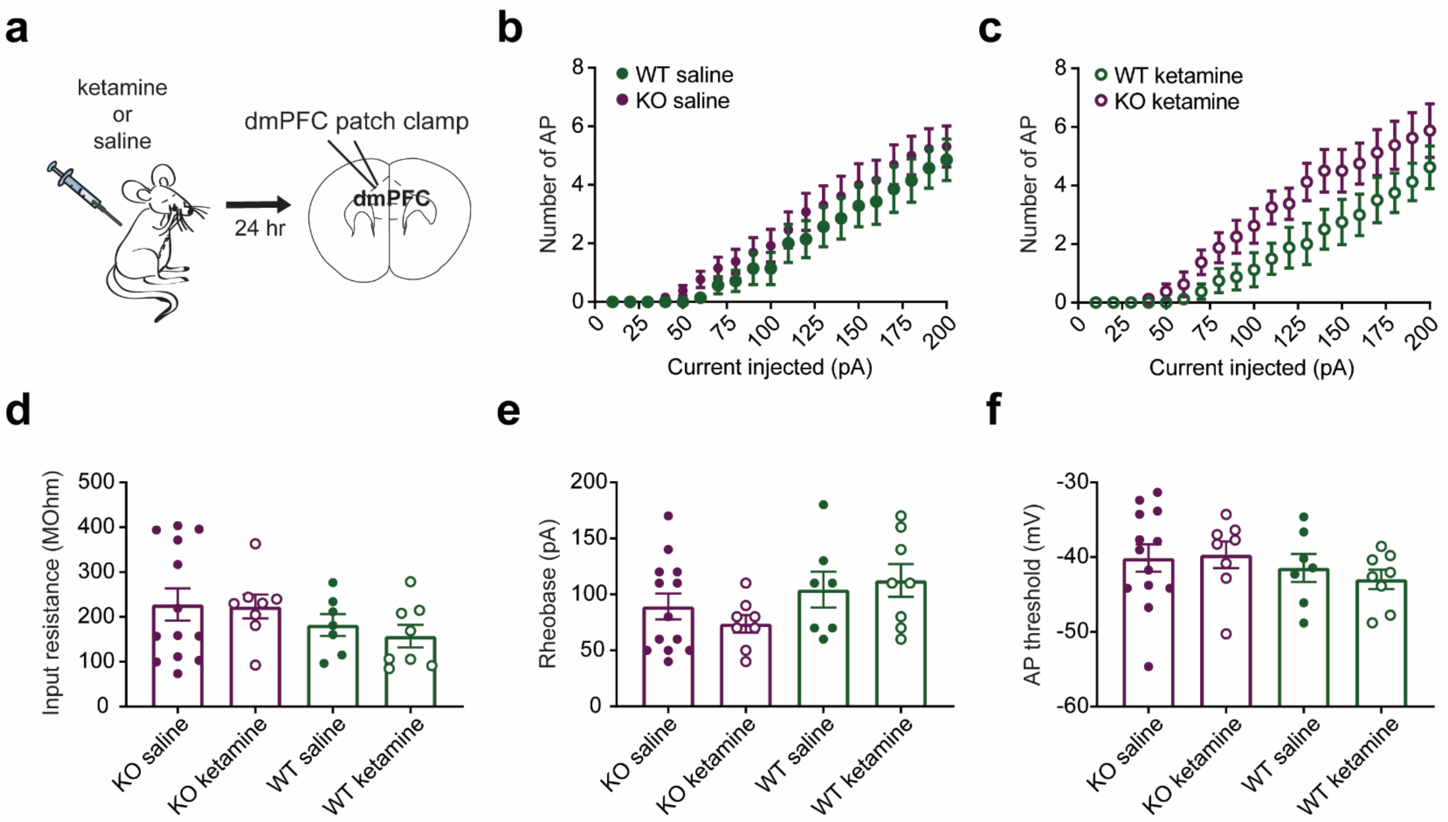


**Supplemental Figure 2. Effects of ketamine and genotype on intrinsic dmPFC cell properties. (a)** Experimental schematic of i.p. injections followed by slice recordings 24 hours later. **(b)** Input-output curves compared between WT saline and KO saline did not reveal any group differences, just a main effect of sequential current injected (Two-way RM ANOVA: interaction $P > 0.05$, current injected $P < 0.0001$, experimental group $P > 0.05$). **(c)** Input-output curves generated from WT ketamine and KO ketamine (30 mg/kg) groups. There is a main effect of current injected and a main interaction between current injected and experimental group, indicating that ketamine could be having different effects on the input-output curves of KOs compared to WTs, but no further effects were seen with multiple comparisons (Two-way RM ANOVA: interaction $P < 0.01$, current injected $P < 0.0001$, experimental group $P > 0.05$). No difference was seen across genotype or treatment condition for **(d)** input resistance (Kruskal-Wallis test $P > 0.05$; WT saline N = 7 cells, KO saline N = 13 cells, WT ketamine N = 8 cells, KO ketamine N = 8 cells) **(e)** rheobase (Kruskal-Wallis test $P > 0.05$) or **(f)** action potential (AP) threshold (Kruskal-Wallis test $P > 0.05$).

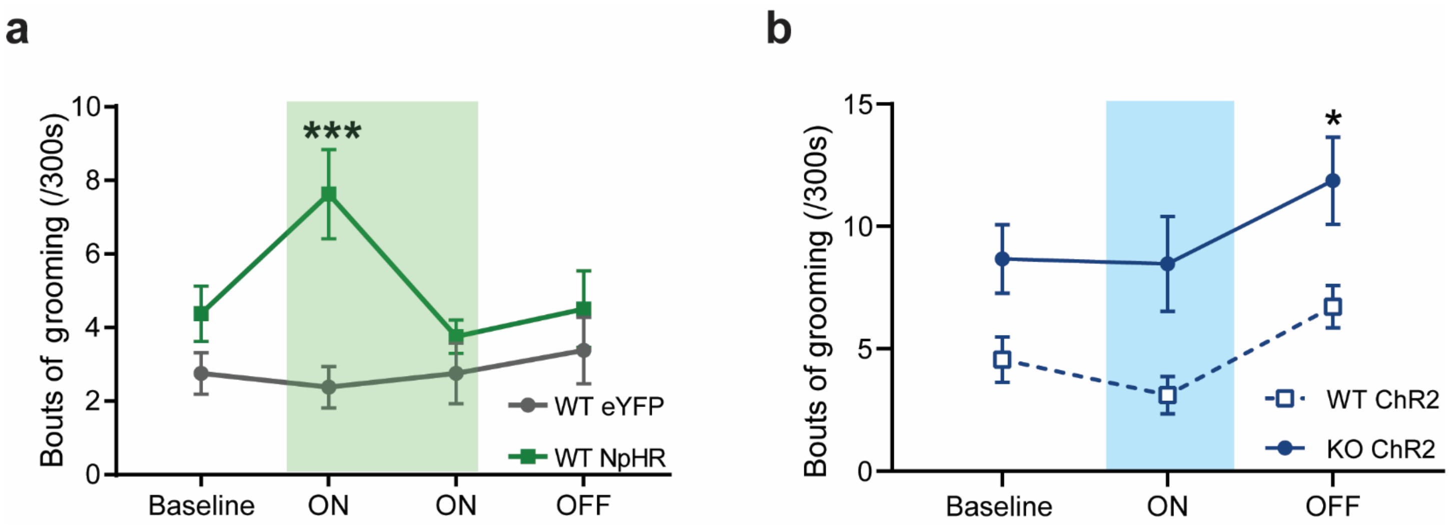


**Supplemental Figure 3. Effects of optogenetic manipulation on frequency of grooming behavior. (a)** Mice expressing NpHR showed increased frequency of grooming during laser on epochs and were significantly different from their eYFP counterparts with a significant main effect of virus and a significant interaction between laser and virus. Multiple comparisons revealed a significant difference between eYFP and NpHR groups during the first 5-minute laser on block (Two-way RM ANOVA: interaction $P<0.05$, laser $P > 0.05$, virus group $P<0.01$; Sidak's multiple comparisons: *** = $P <0.001$). **(b)** SAPAP3 KO mice expressing ChR2 have an elevated frequency of grooming bouts compared to WT mice expressing ChR2 that is not reduced by laser stimulation of dmPFC terminals in the striatum (Two-way RM ANOVA: interaction $P>0.05$, laser $P<0.01$, genotype $P<0.01$; Sidak's multiple comparisons: ** = $P <0.01$).

***Ex-Vivo* Electrophysiology Methods:** To prepare *ex vivo* slices for whole-cell recordings, mice were deeply anesthetized with isofluorane and transcardially perfused with ice-cold glycerol-based slicing solution, decapitated and the brain was removed. Glycerol-based slicing solution contained (in mM) 250 glycerol, 2.5 KCl, 1.2 NaH2PO4, 10 HEPES, 21 NaHCO3, 5 glucose, 2 MgCl2, 2 CaCl2. Coronal slices 250-300 um thick were made on a vibrating microtome (Leica) while the brain was submerged in cold ACSF containing (in mM) 119 NaCl, 26.2 NaHCO3, 2.5 KCl, 1.3 MgSO4, 2.5 CaCl2, 1 NaH2PO4, 11 Glucose; and constantly bubbled with carbogen (95% O2, 5% CO2). From this moment on, slices were constantly submerged in ACSF bubbled with carbogen. They were left in a recovery chamber at 34C for 45-60 minutes, then stored at RT until the recording time. Cells were identified using bright-field, and dmPFC cells were targeted.

For current clamp recordings, patch electrodes (3-6 MOhm tip resistance) were filled with a Potassium-based internal solution containing (in mM): 130 KMeSO3, 8 NaCl, 2 MgCl2, 0.16 CaCl2, 0.5 EGTA, 10 HEPES, 2 MgATP, 0.3 NaGTP, at 290 mOsm, pH 7.2-7.3. To generate APs, neurons in current clamp were depolarized with a series of 20 000-ms current pulses, with 10 pA between each current pulse. This series of current pulses was alternated with hyperpolarizing steps (−50 pA) to examine input resistance.

Electrophysiological recordings and data acquisition were performed with Multiclamp 700B amplifier and pClamp software (Molecular Devices). Analysis was performed with custom-made Excel (Microsoft) macros and pClamp software (Molecular Devices). Holding current and input resistance were continuously monitored as proxies of recording stability.
